# Elevated photic response is followed by a rapid decay and depressed state in ictogenic networks

**DOI:** 10.1101/2022.01.30.478306

**Authors:** Sverre Myren-Svelstad, Ahmed Jamali, Sunniva S. Ophus, Anna M. Ostenrath, Kadir Aytac Mutlu, Helene Homme Hoffshagen, Adriana L. Hotz, Stephan C.F. Neuhauss, Nathalie Jurisch-Yaksi, Emre Yaksi

**Author notes:** These authors contributed equally to this work (Sverre Myren-Svelstad, Ahmed Jamali). Corresponding authors, these authors jointly supervised this work. Corresponding authors.

## Abstract

The switch between non-seizure and seizure states involves profound alterations in network excitability and synchrony. Both increased and decreased excitability may underlie the state transitions, as shown in epilepsy patients and animal models. Inspired by video-electroencephalography recordings in patients, we developed a framework to study spontaneous and photic-evoked neural and locomotor activity in zebrafish larvae. We combined high-throughput behavioral tracking and whole-brain in vivo two-photon calcium imaging to perform side-by-side comparison of multiple zebrafish seizure and epilepsy models. Our setup allowed us to dissect behavioral and physiological features that are divergent or convergent across multiple models. We observed that locomotor and neural activity during interictal and spontaneous ictal periods exhibit great diversity across models. Yet, during photic stimulation, hyperexcitability and rapid response dynamics was well conserved across multiple models, highlighting the reliability of photic-evoked seizure activity for high-throughput assays. Intriguingly, in several models, we observed that the initial elevated photic response is often followed by fast decay of neural activity and a prominent depressed state. We argue that such depressed states are likely due to homeostatic mechanisms triggered by excessive neural activity. An improved understanding of the interplay between elevated and depressed excitability states might suggest tailored epilepsy therapies.

**KEY POINTS:**

- Features of spontaneous locomotor and neural activity varies across zebrafish epilepsy and seizure models.
- We propose photic stimulation as a reliable tool to investigate behavioral and physiological phenotypes in zebrafish epilepsy and seizure models.
- We observed elevated activity with faster dynamics in response to photic stimulation in all tested zebrafish models.
- Photic-evoked neural responses were often followed by depressed state in seizure-prone networks

## INTRODUCTION

Epilepsy is one of the most dynamic brain disorders. The switch between non-seizure and seizure states involves profound alterations in network excitability and synchrony. Increased excitability is often considered as the main seizure generating mechanism (1, 2), and many anti-seizure drugs act by preventing hyperexcitable states (3). Strikingly, decreased excitability is also observed during seizure generation (ictogenesis). For example, hypoactivity of cholinergic neurons in brainstem nuclei and basal forebrain is proposed to induce impaired consciousness in temporal lobe seizures (4, 5). Moreover, enhanced GABAergic inhibition of thalamocortical neurons may underlie absence seizures (6). A wide-spread and transient hypoactivity is the hallmark of post-tonic-clonic-seizure states (7). Consequently, spreading of hypoexcitability across local and global networks may explain important seizure symptoms. Despite such intriguing observations from animal studies and human patients, interplay between hyperexcitable and hypoexcitable states in ictogenic networks remains poorly understood.

Clinical and fundamental applications of systems neuroscience are currently boosting our understanding of epilepsy as a network disorder (8–16). Recording whole-brain activity with high spatiotemporal resolution, however, is challenging in human patients and rodent models due to anatomical and technical constraints. Here, we performed in vivo two-photon calcium imaging across the entire brain of small and transparent zebrafish larvae, to explore the spatial and temporal dynamics of altered network excitability (17–20). Inspired by video-electroencephalography (video-EEG) and photic stimulation protocols that are commonly used in human patients, we performed complementary recordings of animal behavior and whole-brain activity (21). We delivered precise photic stimulation to trigger neural responses, since photosensitivity is an important physiological trait in epilepsy (21–24). Spontaneous locomotor and neural activity displayed a hypoactive or hyperactive phenotype depending on the seizure or epilepsy model (20, 25–30). Photic stimulation evoked increased swim velocities and change in swim angle, in line with tonic-clonic seizure-like behavior observed in zebrafish larvae (26). Intriguingly, photic stimuli induced striking neural dynamics, where an initial elevated state was often followed by a prominent depressed state. Pentylenetetrazole (PTZ), a proconvulsant which is a GABAA antagonist (20, 26, 27), elicited dose-dependent elevated-to-depressed network responses, which was not observed in control animals. The gabra1 knock-out model of the α1 subunit of GABAA receptor (30) resembled low-dose PTZ dynamics.

On the contrary, a genetic model with a dissimilar pathophysiology, the eaat2a knockout model lacking the major astroglial glutamate transporter (29, 31), displayed elevated photic responses followed by less-pronounced depressed states. Taken together, we show that the brain networks’ adaptation to photic stimulation involves dynamic and rapid switching between elevated and depressed states.

## RESULTS

### Spontaneous locomotor and neural activity display a hypoactive or hyperactive phenotype depending on the seizure or epilepsy model

To investigate excitability in pharmacological seizure and genetic epilepsy models, we combined locomotor behavior tracking and two-photon calcium imaging of neural activity. A tonic-clonic-like seizure phenotype in zebrafish larvae consist of an initial phase of high-speed swirl-like swimming followed by a period of immobility (26). Application of the proconvulsant pentylentetrazole (PTZ), a GABAA antagonist, at high dosage (15 mM) led to a clear increase in mean spontaneous swim speed (Fig. 1a, left panel) and change in swim angle (Supplementary Fig. 2, left panel). Lower dosages (1 and 5 mM) did not lead to significant increase in velocities nor swim angle. In gabra1 homozygous mutants with knockout of GABAA receptor α1 subunits, spontaneous swim activity and change in swim angle were not altered (Fig. 1a, Supplementary Fig. 2, center panels). On the contrary, in the eaat2a model lacking the major astroglial glutamate transporter, homozygous mutants were strikingly inactive at baseline, measured both by swim velocity and change in swim angle, in line with a previous study (29) (Fig. 1a, Supplementary Fig. 2, right panels). In the scn1lab knockout model of α1 subunit of NaV1.1 receptors (25, 28, 32), the mean spontaneous swim velocity and change in swim angle was significantly lower compared to controls (Supplementary Fig. 1a, f). Taken together, our findings reveal that the spontaneous locomotor behavior of seizure and epilepsy models may exhibit fundamentally different features ranging from hypo-to hyperactivity.

**Figure 1.**
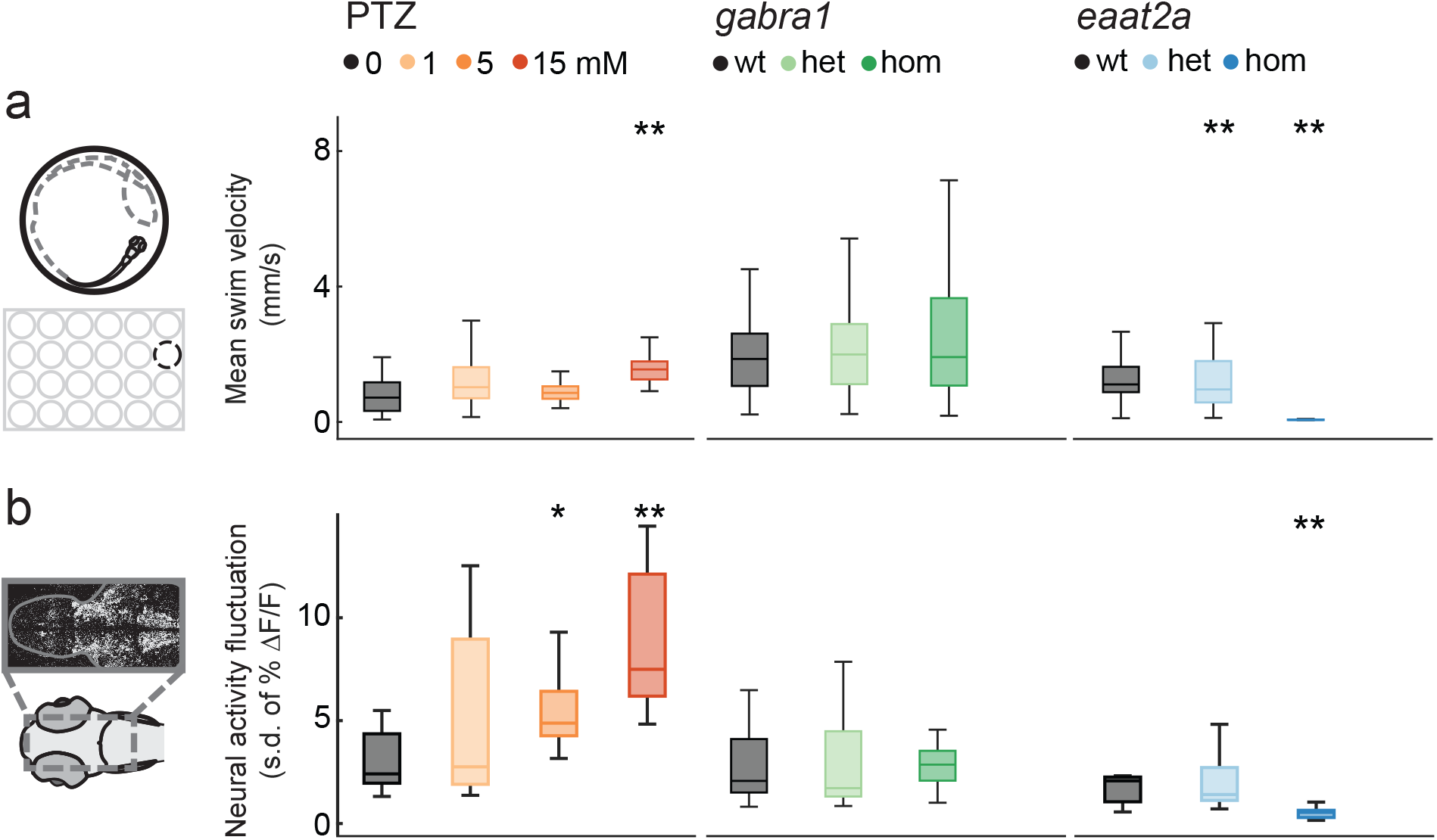
Spontaneous locomotor and neural activity display a hypoactive or hyperactive model-dependent phenotype. **a** Locomotor behavior was recorded in a 24-well plate. Mean swim velocity was calculated per 1 sec time bins during 1-hour baseline recordings in the PTZ, gabra1, and eaat2a models. **b** Neural activity fluctuation was calculated as the standard deviation of ΔF/F (%) during the last 2 min of 1-hour recordings in 5 day old Tg(elavl3:GCaMP6s) zebrafish larvae. Total sample size for behavioral experiments for PTZ (control, n=48; 1 mM, n=48; 5 mM, n=48, 15 mM, n= 48), gabra1 (wildtype, n=39; heterozygous, n=81; homozygous, n=48) and eaat2a (wildtype, n=11; heterozygous, n=41; homozygous, n=42) model. Total sample size for calcium recordings for PTZ (control, n=8; 1 mM, n=7; 5 mM, n=9, 15 mM, n= 10), gabra1 (wildtype, n=9; heterozygous, n=13; homozygous, n=9) and eaat2a (wildtype, n=7; heterozygous, n=13; homozygous, n=11) model. *p<0.05, **p<0.01. Wilcoxon rank-sum test. Boxplots represent median with interquartile ranges, whiskers extend to the most extreme data points that are not outliers.

**Figure 2.**
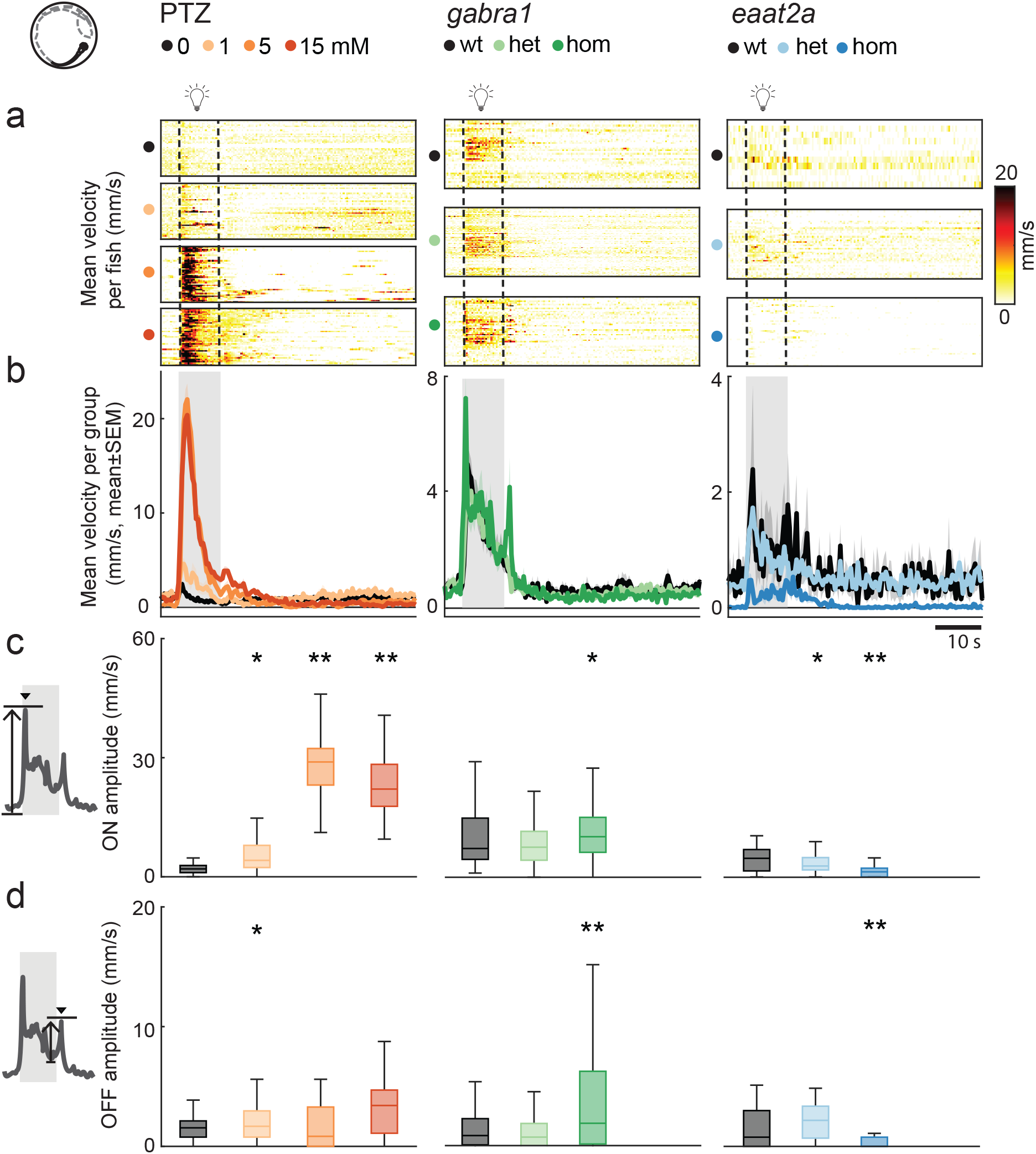
Photic stimulation leads to model-dependent change in swim velocity. **a** Mean swim velocity in response to 10 seconds photic-stimulation with 5-min inter-stimulus interval. Each line in heatmaps represent average across 5 trials for one fish. Dotted lines indicate the start and end of photic stimulation. **b** Mean swim velocity per subgroup. Gray shaded area indicates photic stimulation. **c** Light-on amplitude measured as maximum change in velocity during the first 5 sec after light onset. **d** Light-off amplitude measured as the maximum change in velocity during the first 5 sec after light offset. Total sample size for the PTZ (control, n=42; 1 mM, n=42; 5 mM, n=41, 15 mM, n= 41), gabra1 (wildtype, n=39; heterozygous, n=81; homozygous, n=48) and eaat2a (wildtype, n=11; heterozygous, n=41; homozygous, n=42) model. *p<0.05, **p<0.01. Wilcoxon rank-sum test. Shaded regions represent s.e.m. Boxplots represent median with interquartile ranges, whiskers extend to the most extreme data points that are not outliers.

Next, we examined the neural activity by two-photon microscopy in Tg(elavl3:GCaMP6s) zebrafish larvae expressing calcium indicator GCaMP6s in all neurons. We analyzed the fluctuations (standard deviation) of neural calcium activity during baseline periods (29). The baseline neural activity resembled locomotor behavior in the PTZ, gabra1 and eaat2a models (Fig. 1b). PTZ application led to a dose-dependent increase in calcium signal fluctuations (Fig. 1b, left panel), in line with increased swim velocities observed in the behavioral experiments. In gabra1 homozygous larvae, baseline neural activity was not changed (Fig. 1b, center panel), while homozygous eaat2a animals displayed a significant decrease in baseline neural activity (29) (Fig. 1b, right panel). Analysis of two-photon recordings in the scn1lab homozygous mutants was not possible due to mutation-linked hyperpigmentation. In sum, we observe that locomotor and neural activity at baseline may display a hypoactive or hyperactive phenotype dependent on the model. This underlines that a simplistic readout of epilepsy phenotype purely based on an increase in activity may not encompass the whole pathophysiology of epilepsy.

### Photic stimulation leads to model-dependent change in swim velocity and angle

The occurrence of spontaneous seizures is unpredictable. To assess neural hyperexcitability with a better temporal control, we next used 10 seconds photic stimulation with 5 minutes inter-stimulus-intervals (ISI) in seizure and epilepsy models. By pharmacological perturbation with PTZ, we demonstrated distinct locomotor signatures. We found that the light-on responses were clearly increased for all PTZ concentrations compared to control, with the medium concentration (5 mM) eliciting the strongest mean response (Fig. 2a-c, left panels). We also observed stronger light-off responses in PTZ-treated animals (Fig. 2d). Since previous studies reported swirl-like zebrafish swim-patterns during seizures (26), we next quantified the change in swim angle. Change in swim angle was the largest upon application of higher PTZ dosages (Fig. 3a-d, left panels). The gabra1 homozygous mutants displayed a phenotype similar to higher dose PTZ, with higher amplitude light-on and light-off swim velocity responses (Fig. 2a-d, center panels), but change in swim angle was not significantly different from their controls (Fig. **3a**-d, center panels). The eaat2a homozygous larvae had lower amplitudes both during light-on and light-off responses (Fig. 2a-d, right panels). Swim angle was not altered in eaat2a mutants compared to control (Fig. 3a-d, right panels). In the scn1lab homozygous animals, we observed an increased swim velocity and angle upon light-on compared to wild-type, while light off amplitudes were unaltered (Supplementary Fig. 1b-e, g-j). Taken together, our results using photic stimulation show that photic-evoked swim velocity and change in swim angle are model-dependent.

**Figure 3.**
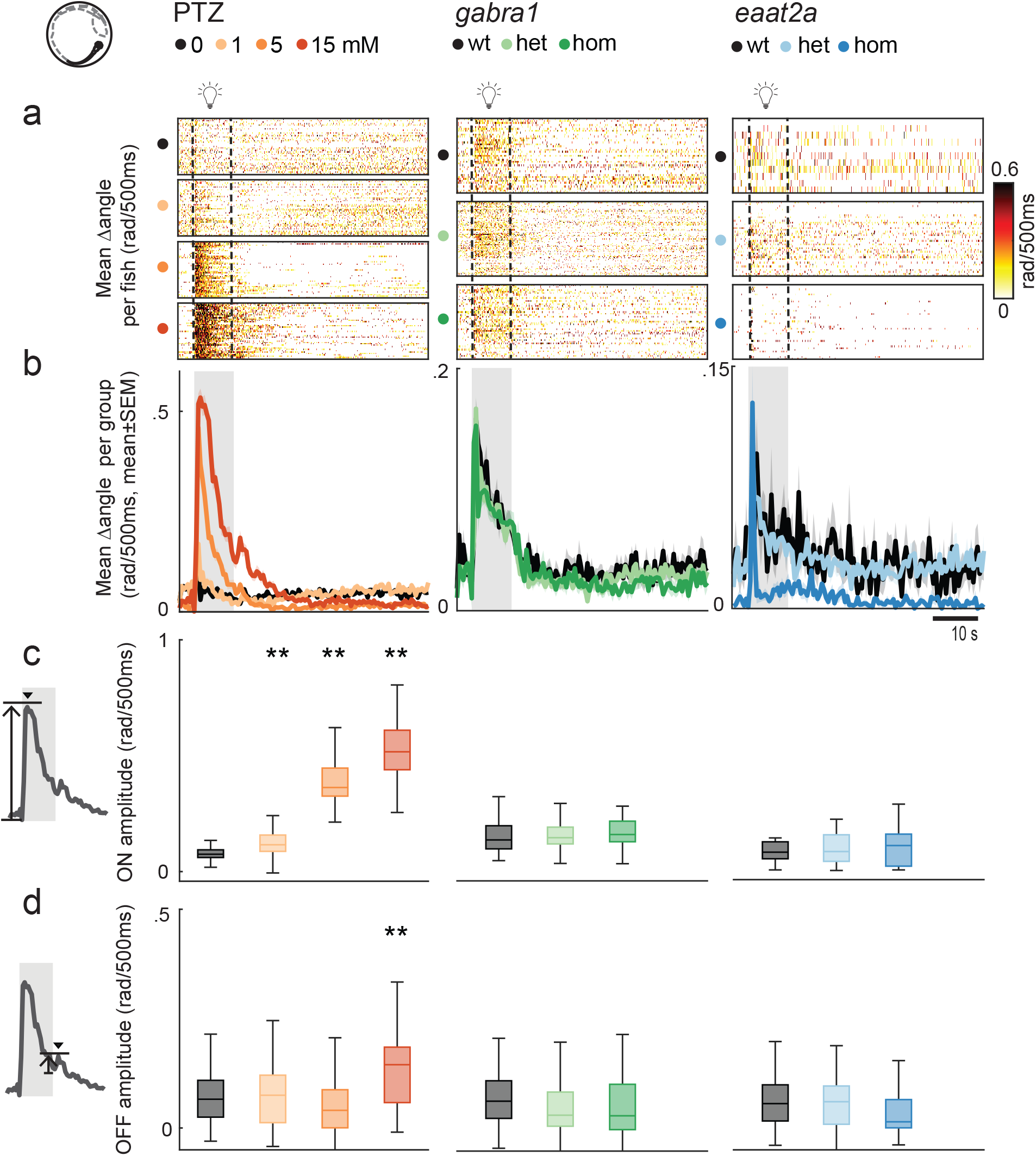
Photic stimulation leads to model-dependent change in swim angle. **a** Mean change in swim angle in response to 10 seconds photic-stimulation with 5-min inter-stimulus interval. Each line in heatmaps represent average across 5 trials for one fish. Dotted lines indicate the start and end of 10 sec photic stimulation. **b** Mean change in swim angle per subgroup. Gray shaded area indicates photic stimulation. **c** Light-on amplitude measured as maximum change in swim angle during first 5 sec after light onset. **d** Light-off amplitude, as the maximum change in swim angle during the first 5 sec after light offset. Total sample size for the PTZ (control, n=42; 1 mM, n=42; 5 mM, n=41, 15 mM, n= 41), gabra1 (wildtype, n=39; heterozygous, n=81; homozygous, n=48) and eaat2a (wildtype, n=11; heterozygous, n=41; homozygous, n=42) model. *p<0.05, **p<0.01. Wilcoxon rank-sum test. Shaded regions represent s.e.m. Boxplots represent median with interquartile ranges, whiskers extend to the most extreme data points that are not outliers.

### Fast dynamics of neural activity and prominent depressed network state are observed following photic stimulation

Next, we studied the strength and dynamics of neural responses across the brain upon 10 seconds photic stimulation with various inter-stimulus-intervals. We calculated the mean photic-evoked response across trials. For the 5-min ISI in the PTZ model, photic responses had significantly shorter latency to peak (5 mM) (Fig. 4a-b, left panel), and higher amplitude (1 and 15 mM) (Fig. 4c, left panel). In homozygous gabra1 mutants, the latency was also shorter, while the amplitude was comparable to controls (Fig. 4a-c, center panels). In homozygous eaat2a larvae, latency was unaltered, but the amplitude was higher (Fig. 4a-c, right panels).

**Figure 4.**
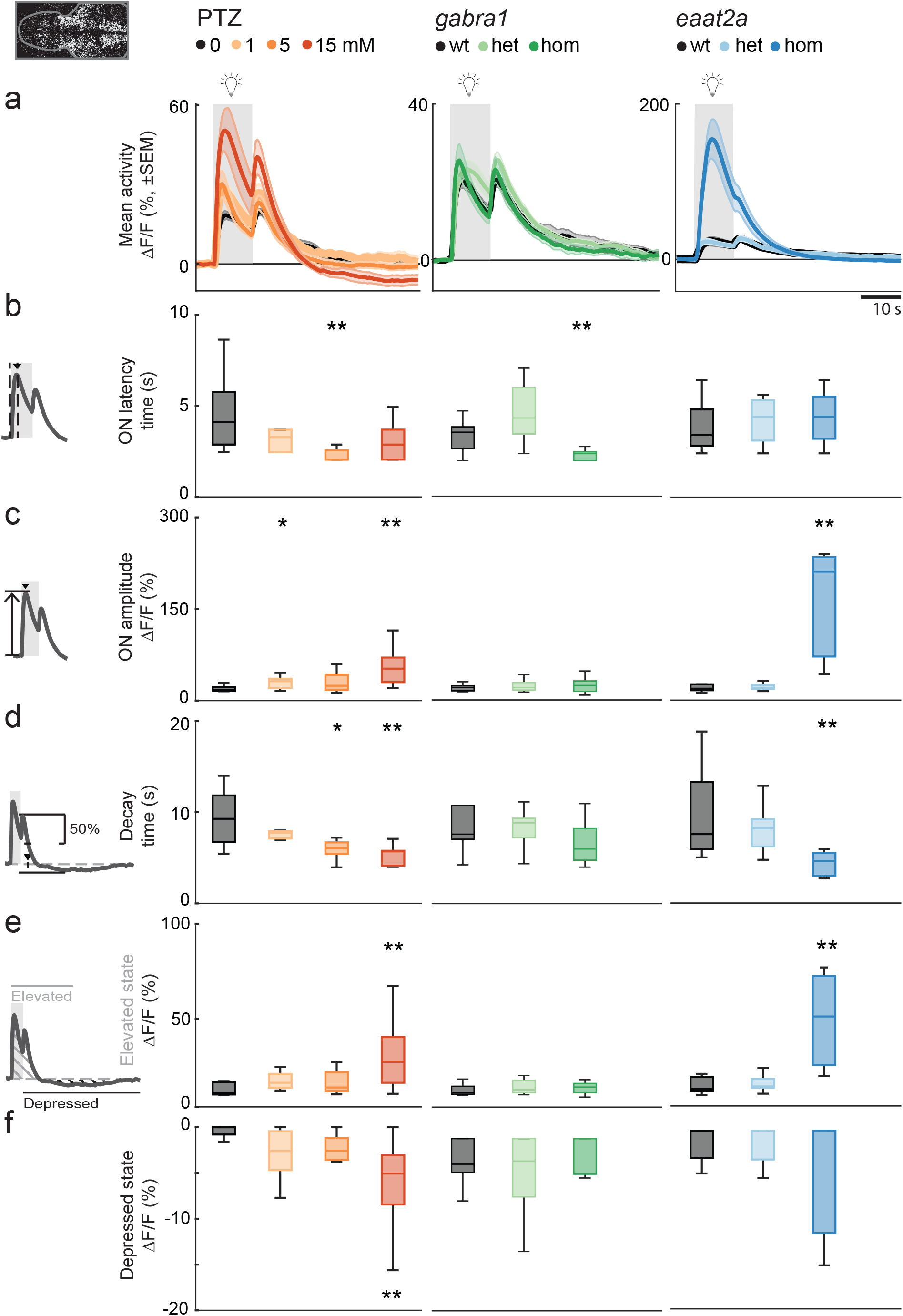
Photic stimulation elicits elevated neural responses with fast dynamics and prominent depressed state. **a** Mean neural calcium responses (ΔF/F, %) to photic stimulation per subgroup averaged across 5 trials of 5-min inter-stimulus-interval. Gray shaded area indicates photic stimulation. **b** Latency to peak amplitude during 10 sec photic stimulation. **c** Peak amplitude during photic stimulation. **d** Time constant for 50% decay of calcium signal from light offset. **e** Elevated state as measured by mean area-under-curve above 2 standard deviations from a 1 min baseline (1 min period after light onset). **f** Depressed state as measured by mean area-over-curve below 2 standard deviations from a 1 min baseline (first 2 minutes after light turned on). Total sample size for the PTZ (control, n=8; 1 mM, n=7; 5 mM, n=9, 15 mM, n= 10), gabra1 (wildtype, n=9; heterozygous, n=13; homozygous, n=9) and eaat2a (wildtype, n=7; heterozygous, n=13; homozygous, n=11) model. For eaat2a homozygous mutants, 4 of 11 animals had only plateau-like events; hence, 7 animals were included in the analyses for this figure. *p<0.05, **p<0.01. Wilcoxon rank-sum test. Shaded regions represent s.e.m. Boxplots represent median with interquartile ranges, whiskers extend to the most extreme data points that are not outliers.

When analyzing dynamics of photic-evoked neural activity, we observed a striking phenomenon: the initial elevated state appeared to be followed by a faster decay and subsequent depressed state (Fig. 4a). To quantify how fast the elevated state decays into a depressed state, we calculated the decay-time constant from the maximum point of the light-off peak. We observed that this faster decay of neural activity is a feature that was preserved across PTZ concentrations, as well as gabra1 and eaat2a models. For 5-min ISI, the differences between control vs 5 or 15 mM PTZ and for homozygous eaat2a were significant, while there was a trend for gabra1 (p=0.16) homozygous larvae (Fig. 4d). Differences were also generally significant for shorter inter-stimulus intervals (Supplementary Fig. 3). This highlights the presence of homeostatic mechanisms that are recruited to quickly respond to elevated neural activity and rapidly reduce it to a less excitable or depressed state.

Next, we quantified these elevated and depressed states by calculating area-under-curve (1 min period from light onset) and area-over-curve (2 min period from light offset), respectively. In the 15 mM PTZ model, elevated and depressed states are correlated (Supplementary Fig. 4), and significantly more prominent compared to control (Fig. 4e-f, left panels). For the gabra1 homozygous larvae, the elevated and depressed states were comparable to control(Fig. 4e-f, center panels). The homozygous eaat2a elevated state was more prominent compared to controls, but we did not observe a significant depressed state on a group level (Fig. 4e-f, right panels).

Occasionally, we also observed photic-evoked events leading to a prolonged elevated state without a following depressed state. These responses displayed dynamics with an elevated ‘plateau’-phase clearly outlasting the photic stimulus, and they were analyzed separately from the briefer and more common ‘non-plateau’ events (Supplementary Fig. 5a-b). For the 15 mM PTZ and gabra1 homozygous animals, ‘plateau-like’ events were very few; only 7/246 and 5/221 events, respectively. On the contrary, a rather high ratio of events in the eaat2a homozygous animals were ‘plateau’-like (42/265), and more than half of the 5-min ISI events (32/55). The ‘plateau’ morphology neural events in eaat2a mutants had very distinct features, with significantly higher amplitudes and slower dynamics compared to ‘non-plateau’ events (Supplementary Fig. 5c-f). This suggests that compensatory mechanisms to balance hyperexcitability are possibly not always as efficiently recruited during prolonged elevated states in this model.

### Photic-evoked state dynamics are brain-region-specific

Next, we asked how the activity of different brain regions are recruited upon photic stimulation. To investigate this, we delineated five brain regions (Fig. 5a, f, k, Fig. 6a) – the telencephalon, thalamus, optic tectum (homologous to superior colliculus in mammals), cerebellum, and brainstem – and analyzed region-wise neural activity. Response dynamics of individual fish (Fig. 5b, g, l), and per subgroup (Fig. 5c, h, m) appeared to be brain-region-dependent.

**Figure 5.**
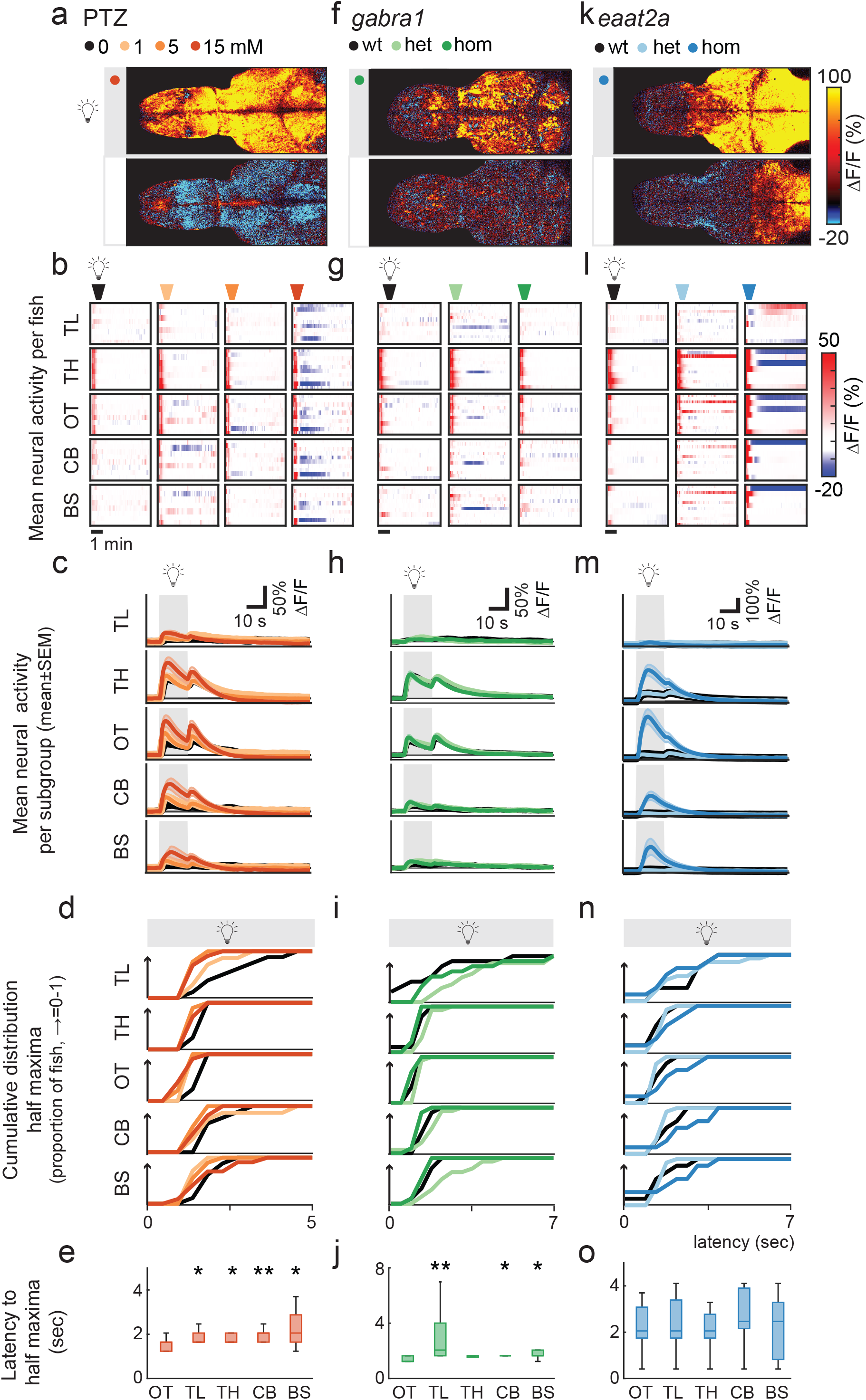
Brain regions are differentially recruited upon photic stimulation. **a, f, k** Representative examples of fish during periods where light was turned on and off. Images show mean calcium signals (ΔF/F) averaged across 5 trials. Upper panels show a time period starting 5 sec after light onset, lower panels from 30 sec after light offset. Both panels show the average activity during a 1.68 second period. **b, g, l** Average calcium signals were extracted from five brain regions: telencephalon (TL), thalamus (TH), optic tectum (OT), cerebellum (CB), and brainstem (BS). Each line in heatmaps represent ΔF/F (%) of one fish averaged across 5 trials of 5-min inter-stimulus-interval. **c, h, m** Average activity per subgroup, ΔF/F (%). **d, i, n** Cumulative distribution of latency to half maxima during photic stimulation. **e, j, o** Latency to half maxima for 15 mM PTZ (e), homozygous gabra1 mutants (j), and homozygous eaat2 mutants (o). Latency for optic tectum is compared with the four other brain regions. Total sample size for the PTZ (control, n=8; 1 mM, n=7; 5 mM, n=9, 15 mM, n= 10), gabra1 (wildtype, n=9; heterozygous, n=13; homozygous, n=9) and eaat2a (wildtype, n=7; heterozygous, n=13; homozygous, n=11) model. For eaat2a homozygous mutants, 4 of 11 animals had only plateau-like events; hence, 7 animals were included in the analyses for this figure. Shaded regions represent s.e.m. *p<0.05, **p<0.01. Wilcoxon rank-sum test. Boxplots represent median with interquartile ranges, whiskers extend to the most extreme data points that are not outliers.

**Figure 6.**
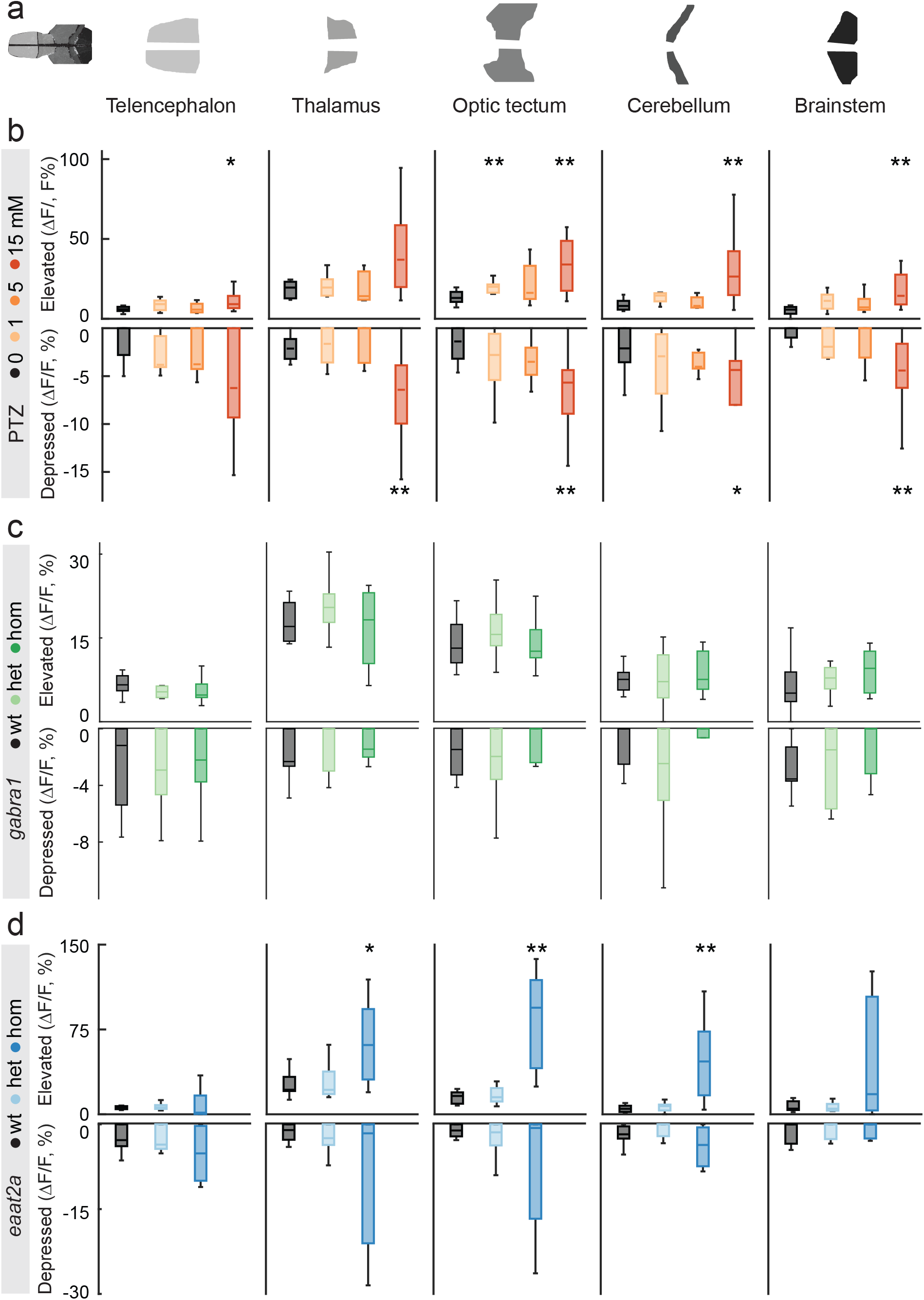
Elevated and depressed states are evoked by photic stimulation. **a** Calculations were done on average calcium signals from separated brain regions: telencephalon, thalamus, optic tectum, cerebellum, and brainstem. **b, c, d – upper panels** Elevated state as measured by area-under-curve above 2 standard deviations from a 1 min baseline (during first minute after light turned on). b, PTZ; c, gabra1; d, eaat2a model. **b, c, d – lower panels** Depressed state as measured by area-over-curve below 2 standard deviations from a 1 min baseline (2 min period from light offset). b, PTZ; c, gabra1; d, eaat2a model. Total sample size for the PTZ (control, n=8; 1 mM, n=7; 5 mM, n=9, 15 mM, n= 10), gabra1 (wildtype, n=9; heterozygous, n=13; homozygous, n=9) and eaat2a (wildtype, n=7; heterozygous, n=13; homozygous, n=11) model. For eaat2a homozygous mutants, 4 of 11 animals had only plateau-like events; hence, 7 animals were included in the analyses for this figure.*p<0.05, **p<0.01. Wilcoxon rank-sum test. Boxplots represent median with interquartile ranges, whiskers extend to the most extreme data points that are not outliers.

To quantify this, we first detected light-on response peak amplitudes to study latency to maximum response of brain regions (Fig. 5d, i, n). We plotted the cumulative proportion of fish in each group which had reached half maximum response across time. A steep rise in the cumulative proportion plot means that the brain region had a short latency to half maximum. Not surprisingly, the neural activity in the optic tectum had the shortest response latency and the strongest amplitude in the PTZ model (Fig. 5c-e). Region-recruitment was significantly more rapid in the optic tectum compared to all other brain regions except from the thalamus in homozygous gabra1 animals (Fig. 5h-j). Recruitment of the optic tectum was not significantly different from other regions in homozygous eaat2a mutants (Fig. 5m-o).

We then explored the elevated and depressed states that we observed across brain regions (Fig. 5 b, g, l, c, h, m), by calculating the area-under-curve above 2 standard deviations from baseline as elevated state, and area-over-curve below 2 standard deviations from baseline as depressed state. For the PTZ seizure model, elevated state was significantly higher for all brain regions except from the thalamus for the 15 mM PTZ group. (Fig. 6b, upper panel). The depressed state was significantly more pronounced for all brain regions except from the telencephalon for the 15 mM group (Fig. 6b, lower panel), and this state was particularly strong in some individuals (Fig. 5b). In gabra1 homozygous larvae, both elevated and depressed states were less-pronounced compared to PTZ-treated larvae, and there were no significant differences between wildtype and mutants (Fig. 6c). In the eaat2a homozygous animals, elevated state was significantly larger for the thalamus, optic tectum, and cerebellum (Fig. 6d, upper panel). While depressed state was similar across subgroups (Fig. 6d, lower panel), a few of eaat2a homozygous animals did display a very distinct depressed state which is seldomly observed in controls (Fig. 5l).

Through systematic analyses of photic-evoked events, we revealed distinct features of neural dynamics across multiple seizure or epilepsy models. Strikingly, an initial elevated state is often followed by a prominent depressed state, which is particularly prominent in the PTZ model. The transition between these two states is rapid in seizure-prone networks. Our findings highlight, that dynamics of ictogenic networks encompass profound and complex alterations of excitability.

## DISCUSSION

Clinical and experimental evidence suggest that seizure generation is a dynamic process consisting of both elevated and depressed network states (4, 33, 34). We show that zebrafish is an efficient model to perform comparative studies on these dynamics in multiple seizure and epilepsy models. The small and transparent zebrafish larvae enable whole-brain two-photon calcium imaging and high-throughput recordings of locomotor readout. First, we demonstrated dose- and model-dependent alterations of spontaneous locomotor and neural activity in line with previous studies (26, 27, 29, 30). Exploration of spontaneous locomotion in multi-well assays is a fruitful screening tool, which has been utilized in large-scale drug screens (26, 27). However, lack of an unbiased metric for seizure-like activity and unpredictable occurrence of seizures may limit the interpretation of such studies. We demonstrated how applying photic stimulation, inspired by human EEG recording paradigms, is a powerful method to perform detailed studies on neural and locomotor dynamics of ictogenesis (23). Our comparative studies on stimulus-locked events enabled us to dissect the dynamic alterations of neural excitability, and to explore whether observed phenomena are generalizable or model-dependent. We propose that our temporally precise photic stimulation approach can enable serial and reliable visualization of neural and astroglial activity as well as neurotransmitters such as glutamate or GABA in future experiments.

We performed in-depth analyses of photic-evoked locomotion during altered excitability states. GABAA-antagonizing perturbations of PTZ and gabra1 models led to increased swim velocities upon photic stimulation. Homozygous scn1lab mutants also showed increased swim velocity compared to controls (23). For the eaat2a knockout model of the major astroglial glutamate transporter, the change in velocity was minor due to general immobility, yet a transition to a mobile state was reproducibly observed upon photic stimulation. Moreover, the change in swim angle was significant in PTZ treated and scn1lab larvae, but not in the gabra1 and eaat2a mutants. This phenotype resembles a commonly described tonic-clonic-like semiology in zebrafish larvae, consisting of high-speed, swirl-like swimming followed by an immobile state (26). The application of photic stimulation enables us to study this phenomenon with precise temporal control.

Next, we analyzed photic-evoked neural activity recorded by two-photon calcium imaging. We explored how the brain circuits evolved during various seizure-inducing perturbations (35). Photic stimulation elicited an initial elevated neural state. The response amplitudes were higher with higher PTZ concentrations and in homozygous gabra1 and eaat2a mutants. The elevated state was most pronounced in the sensory-driven area optic tectum. Propagation of neural hyperexcitability across the entire brain occurred more rapidly in ictogenic networks, especially in animals treated with the GABAA antagonist PTZ. This is in line with the idea that breakdown of surround inhibition underlie seizure spreading (8). Strikingly, we demonstrated that this initial elevated state often was followed by a rapid decay of neural activity and a subsequent depressed state. Interestingly, the elevation of neural activity correlated with depression, especially in the high-dose PTZ model. Our results highlight a fast switching from elevated to depressed state in ictogenic networks.

What might be the underlying mechanisms of rapid switching between elevated and depressed states? It is well-known that elevated (or hyperexcitable) and hypersynchronous brain activity arises during epileptic seizures. Even though depressed states are observed in both patients and animal models, they are less studied and not well understood (5, 34). Landmark studies from the 1960s and 70s in hippocampal and cortical slices revealed the paroxysmal depolarizing shift (PDS) phenomenon (36–39). These electrophysiological recordings were mainly performed in GABAA-antagonist models and demonstrated remarkable neural response dynamics. First, the membrane potential enters a plateau phase with a train of action potential running on top of it, followed by an extended after-hyperpolarization aborting the elevated state. The PDS is hypothesized as the cellular correlate of interictal spikes observed in EEG in patients (36–39). The PDS dynamics are considered to mainly be shaped by ion fluxes through voltage-gated Na+, K+ and Ca2+ channels, and synaptic inputs of excitatory and inhibitory neurons (38). Moreover, astroglia-neuron interactions (20, 29), and effects of neuromodulators like acetylcholine (4) may play important roles. It has been proposed that long lasting plateau phases might be related to seizure propagation (40). Indeed, we observed elevated-to-depressed-state transitions upon photic stimulation in our models, resembling PDS. Moreover, in some of the animals, we even observed long-lasting, elevated plateau-like events clearly outlasting the stimulus. The roles of the depressed (or hyperpolarized) state are enigmatic. Based on the paroxysmal depolarizing shift studies, the depressed states may protect against seizure propagation (39, 41), and cell death due to glutamate excitotoxicity (42). Timing, location, and degree of depressed states may play important roles. The relevance of PDS for human EEG dynamics has yet mostly been inferred from in vitro models. Zebrafish may serve as a novel in vivo model to explore paroxysmal depolarizing shift-like phenomenon in whole-brain networks, which will broaden our understanding of the underlying mechanisms.

While several features of neural dynamics were generalizable across zebrafish seizure and epilepsy models, we also demonstrated differences related to the underlying pathophysiology. Dose-dependent augmentation of elevated state occurred upon treatment with the GABAA antagonist PTZ. However, genetic perturbation of GABAA receptor in gabra1 knockout mutants did, surprisingly, produce less pronounced elevation, perhaps due to genetic compensation during development. Importantly, a previous study reporting photic-induced seizures in gabra1 mutants, was performed in juvenile zebrafish (30), while we examined larval zebrafish which is better suited for high-throughput screens due to ethical constraints. In human patients, gabra1 mutations are linked to juvenile myoclonic epilepsy, where spontaneous and photic reflex seizures typically occur first in youth or young adult age (43). Considering these developmental aspects, it is remarkable that we can detect subtle signs of altered brain excitability already at an early age of larval zebrafish, using photic stimulation. On the contrary, eaat2a knockout of the major astroglial glutamate transporter produced a clearly different response. This model, with decreased uptake of glutamate during synaptic transmission, showed particularly high “light-on” neural response amplitudes, in line with a previous study (29).

Photic stimulation resulted in increased neural and locomotor responses, in line with tonic-clonic seizure-like behavior, in zebrafish larvae. This response may be comparable to photoparoxysmal responses (PPR) in human EEG. Intermittent photic stimulation (IPS) is performed in routine diagnostic EEGs to detect this type of epileptiform activity (21). PPR is seen in several epileptic syndromes including juvenile myoclonic epilepsy (21, 24, 43), and epileptic seizures occur in some patients in association with PPR. EEG responses that outlast the IPS trains, and PPRs not limited to posterior brain regions, are generally considered to have a strong association with epilepsy (21). In our zebrafish seizure models, frequently observed short-lasting elevated photic responses may relate to self-limited PPRs in human patients, while the long-lasting plateau type responses may be comparable to self-sustaining PPRs (44–46).

In our zebrafish model we applied 10 seconds long continuous light stimuli, as this elicited the most reproducible locomotor response. In human patients, flash frequencies between 15 and 20 Hz have been shown to be the most ictogenic. Low-frequency visual evoked potentials (VEP) is mostly limited for testing the functional integrity of the visual system (47), and is less explored in epilepsy. Studies in patients with photosensitive occipital lobe epilepsy reported increased VEP amplitudes consistent with hypersynchrony (48, 49), and abnormal latencies possibly due to failure of cortical gain control (48). VEP may be a valuable tool to investigate brain excitability, especially sub-seizure-threshold network dynamics. We also propose that photic stimulation could be more extensively explored in rodent epilepsy models, where invasive methods like electrical stimulation is most commonly used to induce seizure (22, 50). Further human and animal studies on low-frequency photic stimulation might suggest potential biomarkers in epilepsy and other disorders of altered excitability.

Exploring photic-stimulus-locked events is well-suited for further detailed studies on local and global network dynamics in seizure models. Our findings on depressed network states and seizure propagation may have clear relevance for clinical practice. Traditionally, anti-seizure medications are considered to exert their effects by generally reducing hyperexcitability. More recently, it has been highlighted that different drugs display differential effects on the propagation of the seizure activity (51). We argue that investigating neural mechanism underlying elevated and depressed states in seizure models have great potential. For example, interactions between astroglial and neural networks, or release of neurotransmitters and potassium during photic stimulation could reveal interesting targets to interfere with these states (20). Given all these observations, an improved understanding of the interplay between elevated and depressed excitability states might suggest tailored therapies targeting these states at local and global network level and should be studied further.

## ACKNOWLEDGEMENTS

We thank M. Ahrens (HHMI, Janelia Farm, USA) for the Tg(elavl3:GCaMP6s) transgenic line, P. de Witte (KULeuven, Belgium) for the scn1lab mutant line, and É. Samarut (Université de Montreal, Canada) for the gabra1 mutant line. We thank S. Eggen, V. Nguyen, A.M. Nygaard, and our fish facility support team for technical assistance. We also thank T. Sand, P.M. Omland, and E. Brodtkorb (St. Olav’s University Hospital and NTNU), and the Yaksi lab members for stimulating discussions.

This work was funded by The Liaison Committee for Education, Research and Innovation in Central Norway (‘Samarbeidsorganet’) Grant (S.M.-S., N.J.-Y., E.Y.), Medical Student’s Research Program, NTNU (A.J., S.S.O., H.H.H.), RSO grant, NTNU (A.M.O.), RCN FRIPRO Research Grant 239973 (E.Y.) and ERC starting grant 335561 (E.Y.). Work in the E.Y. lab is funded by the Kavli Institute for Systems Neuroscience at NTNU.

## AUTHOR CONTRIBUTIONS

Conceptualization, A.J. and N.J.-Y.; Methodology, A.J., S.M.-S., N.J.-Y., E.Y.; Data Analysis, S.M.-S., A.J.; Software, S.M.-S., A.J., A.M.O., K.A.M.; Providing reagents and data, A.L.H., S.C.F.N.; Investigation, all authors; Writing, S.M.-S., A.J., N.J-Y., E.Y.; Review & Editing, all authors; Funding Acquisition, S.M.-S., N.J.-Y., E.Y.; Supervision, N.J.-Y., E.Y.

## MATERIALS AND METHODS

### Contact for Reagent and Resource Sharing

Further information and requests for reagents may be directed to and will be fulfilled by the lead author Emre Yaksi.

### Experimental Model and Subject Details

#### Fish maintenance

Fish were kept in 3.5-liter tanks at a density of 15-20 fish per tank in a Techniplast Zebtech Multilinkinsystem at constant conditions: temperature 28 °C, pH 7, 6.0 ppm O2 and 700 μS, at a 14:10 hour light/dark cycle to simulate optimal natural breeding conditions. Fish received a normal diet of dry food (Zebrafeed, Sparos I&D Nutrition in Aquaculture, <100-600, according to their size) two times per day and Artemia nauplii once a day (Grade0, platinum Label, Argent Laboratories, Redmond, USA). Larvae were maintained in egg water (1.2 g marine salt, 20 L RO water, 1:1000 0.1% methylene blue) from fertilization to 3 days post fertilization (dpf) and in artificial fish water (AFW, 1.2 g marine salt in 20 L RO water) from 3 to 5 dpf. The animal facilities and maintenance of the zebrafish, Danio rerio, were approved by the Norwegian Food Safety Authority (NFSA). All experimental procedures were performed on zebrafish larvae up to 5 dpf and in accordance with the directive 2010/63/EU of the European Parliament and the Council of the European Union and the Norwegian Food Safety Authorities.

For experiments, the following fish lines were used: Tg(elavl3:GCaMP6s) (20, 52), eaat2a (29), gabra1udm103 (30), scn1labs552 (25) and wild-type.

#### Behavior tracking and experimental design

Behavior tracking was performed on 5 dpf wildtype (n=216 for 3-hour; n=168 for 1-hour PTZ experiments), gabra1 (n=168), eaat2a (n=120), and scn1lab (n=264) larvae. We used a commercially available automated tracking system (Zantiks inc.), which has a built-in tracking software that filters out “non-realistic” swimming epochs with threshold of 5 pixels, and control of light stimuli. Animals were placed in separate wells of a 24-well cell culture dish. For the PTZ experiments, wells were filled with either 2 mL of PTZ or AFW, according to experimental subgroup. For the other experiments, all wells were filled with 2 mL AFW. Fish were placed into the wells right before the start of the experiments. After 1 hour of tracking under light conditions for acclimatization and baseline readout, the lights were turned off for 10min, then followed by “white” light stimulation of 10 sec durations applied with inter-stimulus intervals (ISI) of 10, 5, 2, and 1 min. Total recording time was 3 hr. Because long recording length leads to a high total drug exposure, we additionally performed shorter 1-hour recordings for the PTZ model. Here, we used a 10 min baseline (in darkness), followed by 10 sec light stimuli with ISI 5, 2, 1, 0.5, and 0.25 minute.

#### Two-photon calcium imaging

Two-photon calcium imaging was performed on 5 dpf Tg(elavl3:GCaMP6s) (n=41, for PTZ experiments), gabra1;Tg(elavl3:GCaMP6s) (n=31), and eaat2a;Tg(elavl3:GCaMP6s) (n=38) zebrafish larvae. Animals were paralyzed upon injection of α-bungarotoxin (Invitrogen BI601, 1 mg/mL) (20, 53) and embedded in 1.5% LMP agarose in a recording chamber (Fluorodish, World Precision Instruments). After 10 min of agarose solidification either 0.75 mL (for PTZ experiments) or 1.5 mL (for the genetic models) AFW was added on top of the agarose. Right before start of PTZ experiments, 0.75 mL of either 2, 10 or 30 mM PTZ was added to the AFW, for final concentrations of 1, 5 or 15 mm PTZ. The recordings were performed in a two-photon microscope (Scientifica Inc) using a 16x water immersion objective (Nikon, NA 0.8, LWD 3.0, plan). A mode-locked Ti:Sapphire laser (MaiTai Spectra-Physics) tuned to 920 nm was used for excitation. Volumetric recordings of ten planes of 1536 x 650 pixels were obtained using a Piezo element with an acquisition rate of 2.43 Hz. For a subset of the recordings, single plane recordings of 1536×650 pixels were obtained with an acquisition rate of 24.3 Hz and down-sampled to 2.43 Hz before analysis. Spontaneous calcium activity was recorded for ten minutes in darkness, followed by a stimulus train of 10 seconds long red light using an LED (LZ1-00R105, LedEngin; 625-nm) driven by a custom controlled Arduino device (54). Inter stimulus intervals were 5, 2-, 1-, 0.5-, and 0.25-min. Total duration of the recordings were 60 minutes. The cerebral blood flow was assessed in the prosencephalic arteries or anterior cerebral veins before and after the experiment, and animals without cerebral blood flow were excluded from the final analysis.

#### Genotyping

Following the experiments, larvae from mutant lines were kept for genotyping. For eaat2a and gabra1 lines, genotypes were determined by real-time qPCR melt-curve analyses(29, 30). For scn1lab genotyping, KASP assay (LGC Biosearch Technologies) was performed (55).

### Data analysis

Behavioral data from the tracking system was retrieved as .csv files containing information on swimming distance, XY position over time, and timing of photic stimulation. Custom made MATLAB scripts were used to process and analyze the data. Fish with extreme velocities were flagged as possible tracking error using a script assessing 1-second-binned velocities over the whole recording. The thresholds for flagging were calculated as the 99.96th percentile of the overall-pooled velocities for each model. Subsequently, videos of the experiments with flagged time periods were visually assessed for tracking errors, and afterwards the fish were excluded if tracking errors were found.

Spontaneous behavioral activity was assessed as the mean velocity over the baseline period (1st hour of the 3-hour long experiments). The velocities were compared within subgroups of each model using a Wilcoxon rank sum test (MATLAB built-in function) with a significance level of 0.05. For calculating mean photic-evoked swim velocity, we first averaged the velocity per five trials for each fish; and then analyzed the 5-min ISI stimulations. For detection of light-on and -off peak amplitudes, we found the maximum change in velocity during the first 5 sec after lights were turned on and first 5 sec after the lights were turned off, respectively. Change in swim angle was calculated using MATLAB-inbuilt functions. From three subsequent points of the tracking, two vectors were defined. Using the “det” function, the determinant of the vectors was calculated, and the scalar product was produced using the “dot” function. The angle was subsequently calculated as the arctangent of the determinant and the dot product using “atan2”.

Two-photon microscopy images were aligned using an adapted algorithm (53, 54) that correct for occasional drift in the XY dimension based on hierarchical model-based motion estimation. Regions of interest (ROIs) were manually drawn on one single plane of maximal information. A total of 10 ROIs were drawn; left and right hemispheric ROIs of the following brain regions: telencephalon, thalamus, optic tectum, cerebellum, and brainstem. For each ROI, the fractional change in fluorescence (ΔF/F) relative to a baseline was calculated (56). For spontaneous activity calculation, the last 2 min of the 1-hour recording was selected for calculation. For intermittent photic stimulation, the 5 sec fluorescence preceding the stimuli was the baseline. Linear regression of elevated vs depressed state was calculated in Matlab using “fitlm” function.

### Quantification and Statistical Analysis

Statistical analysis was done using MATLAB. Wilcoxon rank sum test was used for non-paired analysis. P<0.05 was considered as statistically significant.

### Data Availability

The datasets supporting the current study have not been deposited in a public repository but are available from the corresponding author upon request.

### Code Availability

The codes supporting the current study have not been deposited in a public repository but are available from the corresponding author upon request.

