## Supplementary Figures for "Elevated photic response is followed by a rapid decay and depressed state in ictogenic networks"

### Supplementary Material

#### Supplementary Fig. 1

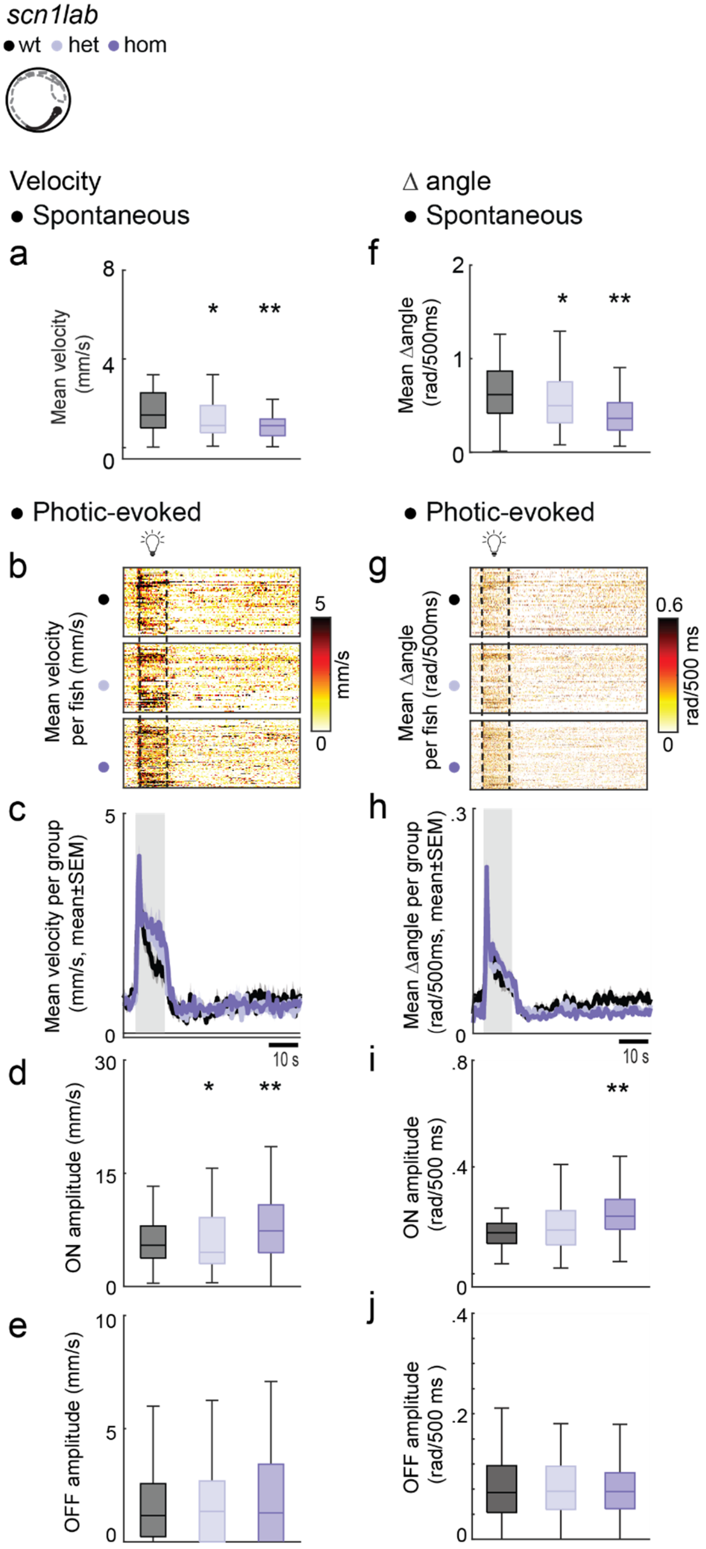


*scn1lab* mutants exhibit hypoactivity during spontaneous locomotion and hyperactivity during photic stimulation

**a** Mean spontaneous swim velocity was calculated per 1 sec time bins during 1-hour baseline recordings. **b** Mean photic-evoked swim velocity for 5-min inter-stimulus interval. Each line in heatmaps represent average across 5 trials for one fish. Dotted lines indicate 10 sec photic stimulation. **c** Mean swim velocity per subgroup. Gray shaded area indicates photic stimulation. **d** Light-on amplitude measured as maximum velocity during 5 sec after light onset. **e** Light-off amplitude, as the maximum velocity during the first 5 sec after light offset. **f** Mean change in swim angle during 1-hour baseline recordings. **g** Mean change in swim angle for 5-min inter-stimulus interval. Each line in heatmaps represent average across 5 trials for one fish. Dotted lines indicate 10 sec photic stimulation. **h** Mean change in swim angle per subgroup. Gray shaded area indicates photic stimulation. **i** Light-on amplitude measured as maximum change in swim angle during 5 sec after light onset. **j** Light-off amplitude, as the maximum change in swim angle during the first 5 sec after light is turned off.
Total sample sizes for *scn1lab* (n=264) larvae (wildtype, n=56; heterozygous, n=75; homozygous, n=125).
*p<0.05, **p<0.01. Wilcoxon rank-sum test. Boxplots represent median with interquartile ranges, whiskers extend to the most extreme data points that are not outliers.

Supplementary Fig. 2. Related to Fig. 1

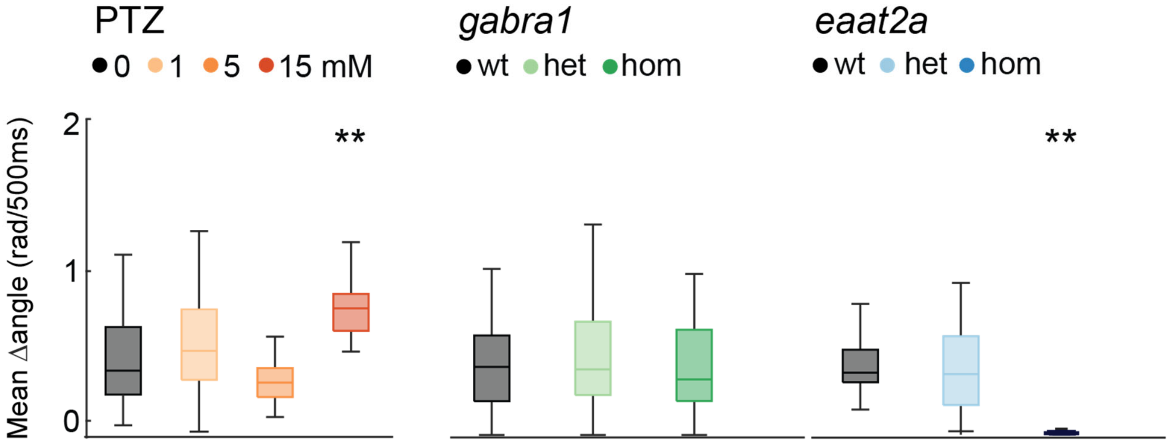


Change in spontaneous swim angle is model-dependent
Mean change in swim angle during 1-hour baseline recordings in the PTZ (left panel), *gabra1* (center panel), and *eaat2a* (right panel) models.
Total sample size for PTZ (control, n=48; 1 mM, n=48; 5 mM, n=48, 15 mM, n= 48), *gabra1* (wildtype, n=39; heterozygous, n=81; homozygous, n=48) and *eaat2a* (wildtype, n=11; heterozygous, n=41; homozygous, n=42) model.
*p<0.05, **p<0.01. Wilcoxon rank-sum test. Boxplots represent median with interquartile ranges, whiskers extend to the most extreme data points that are not outliers.

Supplementary Fig. 3. Related to Fig. 4

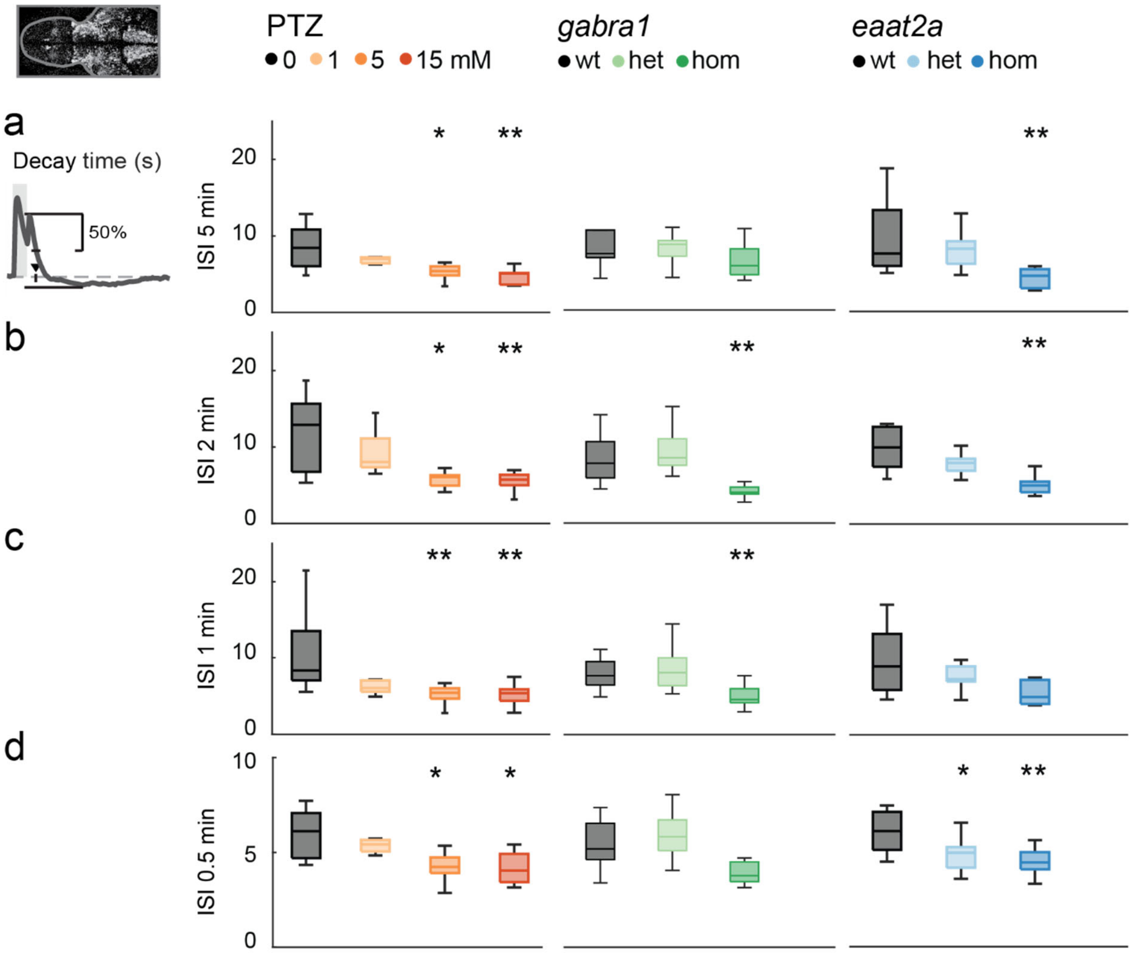


Rapid decay of neural activity is observed after photic stimulation in PTZ, *gabra1* and *eaat2a* models
Time constant for 50% decay of calcium signal from light offset peak for the PTZ (left panels), *gabra1* (center panels) and *eaat2a* models (right panels). **a** Inter-stimulus-interval (ISI) 5-min. **b** ISI 2-min. **c** ISI 1-min. **d** ISI 0.5-min.
Total sample size for the PTZ (control, n=8 ; 1 mM, n=7; 5 mM, n=9, 15 mM, n= 10), *gabra1* (wildtype, n=9; heterozygous, n=13; homozygous, n=9) and *eaat2a* (wildtype, n=7; heterozygous, n=13; homozygous, n=11) model. For *eaat2a* homozygous mutants, 4 of 11 animals had only plateau-like events; hence, 7 animals were included in the analyses for this figure.
*$p<0.05$, **$p<0.01$. Wilcoxon rank-sum test. Boxplots represent median with interquartile ranges, whiskers extend to the most extreme data points that are not outliers.

Supplementary Fig. 4. Related to Fig. 4

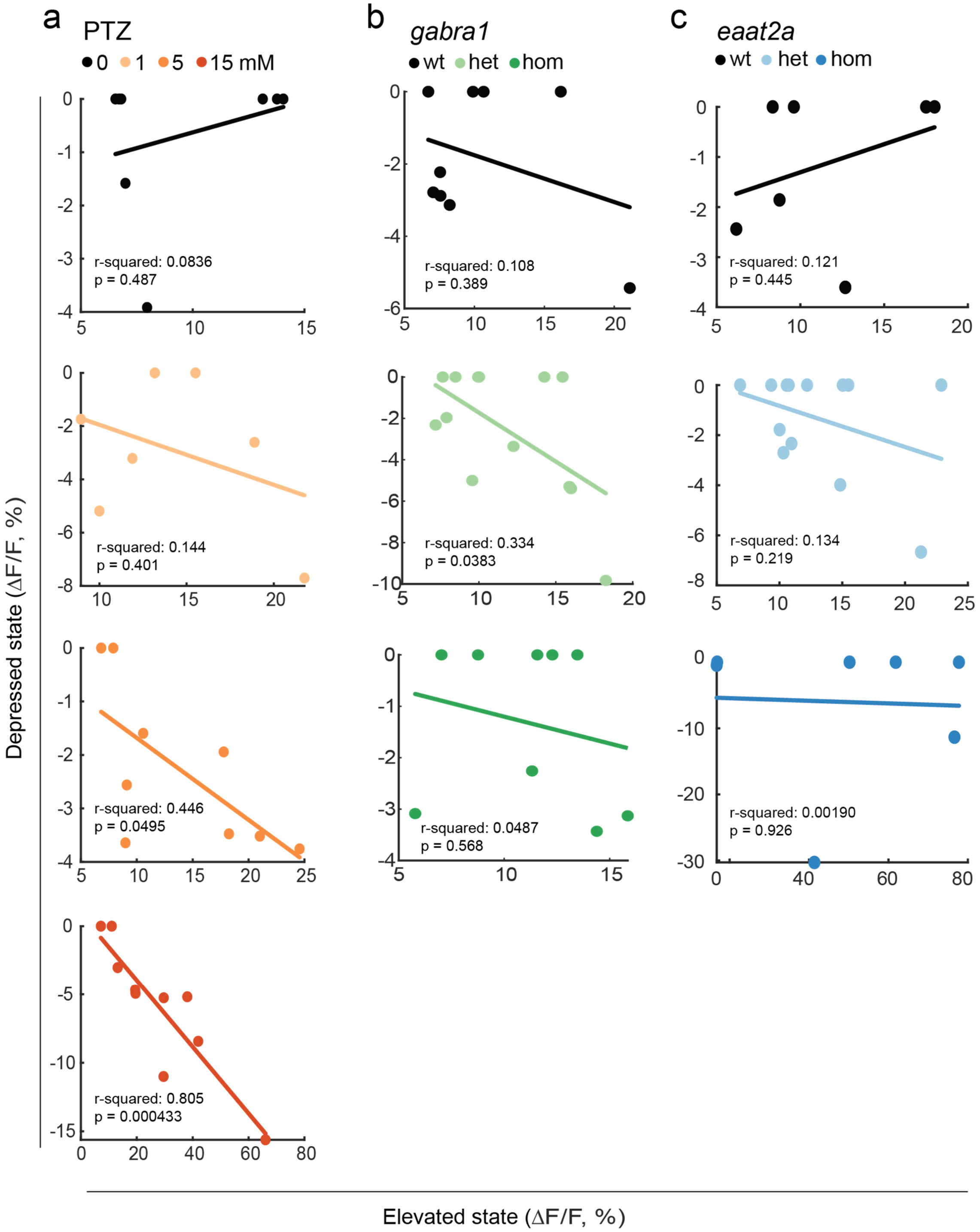


Elevated and depressed states are highly correlated in high-dose pentylentetrazole model
**a** Correlation between elevated and depressed neural state in the PTZ model.
Elevated state as measured by area-under-curve above 2 standard deviations from a 1 min baseline (during first minute after light turned on). Depressed state as measured by area-over-curve below 2 standard deviations from a 1 min baseline (2 min period from light offset).
Both states are given as average ΔF/F (%). **b** Correlation in the *gabra1* model. **c** Correlation in the *eaat2a* model.

Total sample size for the PTZ (control, n=8 ; 1 mM, n=7; 5 mM, n=9, 15 mM, n= 10), *gabra1* (wildtype, n=9; heterozygous, n=13; homozygous, n=9) and *eaat2a* (wildtype, n=7; heterozygous, n=13; homozygous, n=11) model. For *eaat2a* homozygous mutants, 4 of 11 animals had only plateau-like events; hence, 7 animals were included in the analyses for this figure.

#### Supplementary Fig. 5. Related to Fig. 4

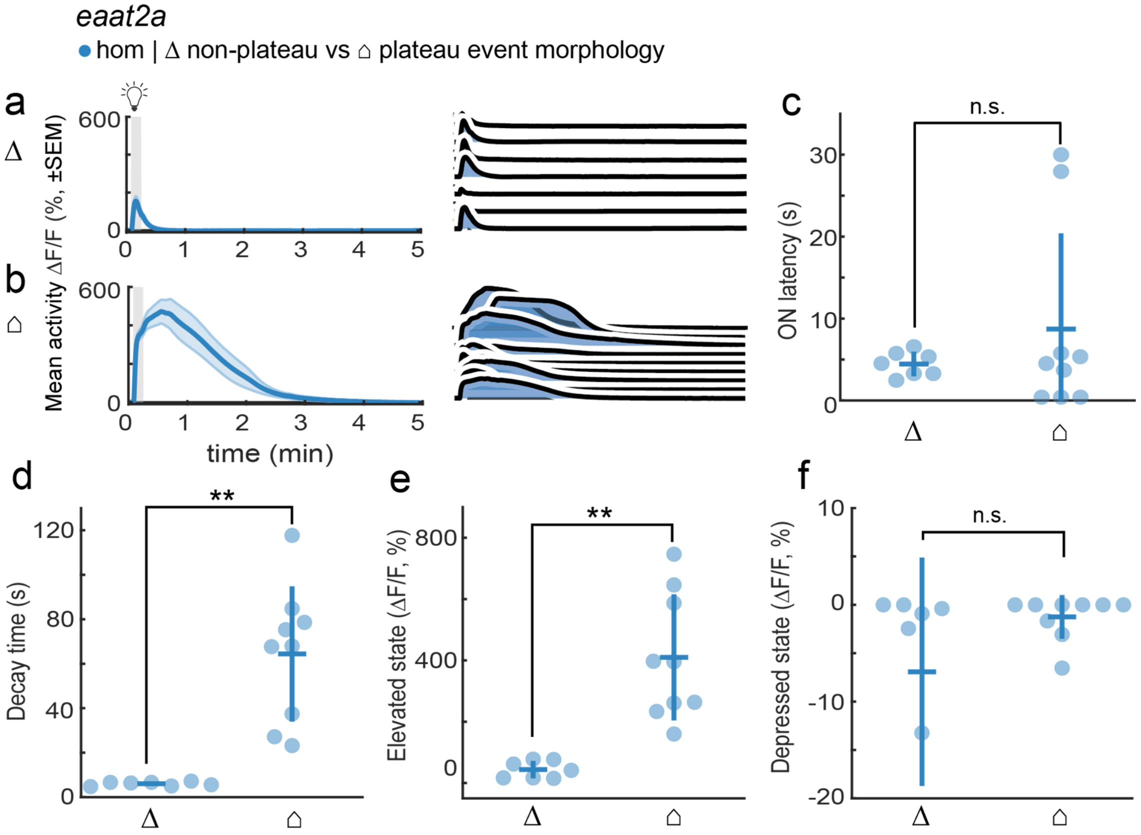


Photic-evoked neural activity in homozygous *eaat2a* animals displays non-plateau or plateau morphology

**a-b** Mean photic-evoked ΔF/F (%) of neural regions averaged across 5 trials of 5-min inter-stimulus-interval. Gray shaded area indicates photic stimulation. Non-plateau events are brief and with a clear peak (**a**), while plateau events have a long duration (**b**). Left panels show activity across group (mean±s.e.m.). Right panels show activity per fish. **c** Latency to ON peak amplitude. **d** Time constant for 50% decay of calcium signal from light offset. **e** Elevated state as measured by mean area-under-curve above 2 standard deviations from a 1 min baseline (during 1 min period after light onset). **f** Depressed state as measured by mean area-over-curve below 2 standard deviations from a 1 min baseline (during a 2 min time period from light offset).

A total of 11 *eaat2a* homozygous mutants were analyzed. For 5-min ISI photic stimulation, two animals had only non-plateau events. Four animals had only plateau events. Five had both non-plateau and plateau events.

*p<0.05, **p<0.01. Wilcoxon rank-sum test. Each data point in scatter plots represent individual fish, error bars represent mean±s.d. per subgroup. Shaded regions represent s.e.m.
